# RecA-NT homology motif in ImuB is essential for mycobacterial ImuA’-ImuB protein interaction and mutasome function

**DOI:** 10.1101/2023.03.28.534377

**Authors:** Joana A. Santos, Kęstutis Timinskas, Meindert H. Lamers, Česlovas Venclovas, Digby F. Warner, Sophia J. Gessner

## Abstract

The mycobacterial mutasome – minimally comprising ImuA’, ImuB, and DnaE2 proteins – has been implicated in DNA damage-induced mutagenesis in *Mycobacterium tuberculosis*. ImuB, predicted to enable mutasome function via its interaction with the β clamp, is a catalytically inactive member of the Y-family of DNA polymerases. Like other members of the Y family, ImuB features a recently identified amino acid motif with homology to the RecA-N-terminus (RecA-NT). In RecA, the motif mediates oligomerization of RecA monomers into RecA filaments. Given the role of ImuB in the mycobacterial mutasome, we hypothesized that the ImuB RecA-NT motif might mediate its interaction with ImuA’, a RecA homolog of unknown function. To investigate this possibility, we constructed a panel of *imuB* alleles in which RecA-NT was removed, or mutated. Results from microbiological and biochemical assays indicate that RecA-NT is critical for the interaction of ImuB with ImuA’. A region downstream of RecA-NT (ImuB-C) also appears to stabilize the ImuB-ImuA’ interaction, but its removal does not prevent complex formation. In contrast, replacing two key hydrophobic residues of RecA-NT, L378 and V383, is sufficient to disrupt ImuA’-ImuB interaction. To our knowledge, this constitutes the first experimental evidence showing the role of the RecA-NT motif in mediating the interaction between a Y-family member and a RecA homolog.

## INTRODUCTION

The mycobacterial mutasome, which has been implicated in DNA-damage tolerance and induced mutagenesis, is minimally composed of a RecA homolog (ImuA’), a catalytically inactive Y-family DNA polymerase (ImuB) and an error-prone C-family DNA polymerase (DnaE2)^1–4^. Although experimental evidence supports the role of DnaE2 as error-prone translesion synthesis (TLS) polymerase^1^, the functions of the ImuA’ and ImuB accessory proteins remain elusive. ImuB lacks the acidic amino acids required for DNA polymerase activity but contains a functional β clamp-binding motif that is thought to mediate access of DnaE2 (which lacks a β clamp-binding motif) to DNA. In support of this hypothesis, mutations in the ImuB β clamp-binding motif abolish DNA-damage induced mutagenesis^2^ and abrogate ImuB-β co-localisation *in vivo*^5^; however, the precise molecular function of ImuB is not known. The function of ImuA’ is even less clear, but homology to RecA makes it tempting to postulate a structural role for ImuA’ similar to that of RecA in binding components of a mutagenic complex and regulating its activity^2,6,7^. Notably, recent work has implicated the myxococcal ImuA protein in inhibition of recombination repair by directly binding RecA, thereby facilitating a switch from error-free repair to TLS^8^.

The central role of ImuB can be ascribed to its multi-protein binding capacity, which is facilitated by its C-terminal region. A yeast two-hybrid assay suggested that ImuB can interact with ImuA’, DnaE2, and itself, as well as the *dnaE1*-encoded high fidelity replicative DNA polymerase^2^. An attempt in that same study to identify an ImuB-interaction region at the N-terminus (first 48 residues) of ImuA’ failed, while a C-terminal deletion (40 residues) seemed to abolish interaction with ImuB. However, one major limitation of the yeast two-hybrid system is the inability to rule out the loss of structural integrity consequent on the engineered mutant alleles as a cause of failed interactions^2^.

Recently, a motif was identified in the C-terminal regions of some active and inactive Y-family polymerases that resembles the N-terminal region of *E. coli* RecA (RecA-NT) responsible for oligomerization^7^. This RecA-NT motif enables binding to the neighbouring RecA molecule in the filament, facilitating oligomerization. Both structural predictions and experimental evidence suggest that the RecA-NT motif mediates the interaction between RecA and UmuC – an essential subunit of the paradigmatic Y-family TLS polymerase, *E. coli* DNA Polymerase V – and that this interaction might be ubiquitous, including even catalytically inactive Y-family members such as ImuB. Furthermore, given the structural similarity between ImuA’ and RecA, it might be expected that the RecA-NT motif in ImuB binds to ImuA’.

Here, we used computational modelling to derive a putative structure of the complex between mycobacterial ImuA’ and ImuB proteins and to evaluate potential features underlying their interaction. Based on the model, we identified regions and specific residues in the RecA-NT motif predicted to be key in maintaining a stable interaction between ImuA’ and ImuB. Using this information, we designed mutations to disrupt the interaction interface, then assessed their impact *in vitro* in biochemical assays and *in vivo* in complementation experiments utilizing a Δ*imuA*’Δ*imuB* double deletion mutant of *Mycobacterium smegmatis*, a mycobacterial model organism^2^. Our results indicate that RecA-NT is essential to secure a stable ImuA’-ImuB interaction and, consequently, mutasome function. Truncation of the RecA-NT motif or substitution of a pair of key hydrophobic residues at the predicted ImuA’-ImuB interface abolishes mutasome function. To our knowledge, this is the first report showing the importance of RecA-NT in binding a RecA homolog.

## MATERIALS AND METHODS

### Computational modelling and analysis

For the initial structural model of the ImuA’-ImuB interaction, the HHpred server^9^ was used to identify modelling templates and to generate sequence-structure alignments. The selected best template corresponded to the *Escherichia coli* RecA-ssDNA filament (PDB id: 3CMU). A homology model was generated using Modeller^10^. Only the RecA-NT-like motif of ImuB (NCBI id ABK76097.1, residues 360-387) and the core of ImuA’ (NCBI id ABK74665.1, residues 63-194) were modelled.

Subsequently, a structural model of the complex formed by the full-length ImuA’ (NCBI id ABK74665.1) and ImuB (NCBI id ABK76097.1) was generated using a local installation of AlphaFoldMultimer v2^11,12^ with default parameters. The ImuA’ N-terminus (residues 1-64) in the model appeared unstructured and was removed from subsequent analyses.

All structural analyses and visualizations were performed using UCSF Chimera^13^. Residue-residue contacts at the interaction interface were analysed using VoroContacts^14^.

### Bacterial strains and growth conditions

All *E. coli* strains were grown in LB culture medium at 37°C with shaking, with the addition of kanamycin (50 µg/ml) where appropriate. All *M. smegmatis* strains were cultured in standard Difco™ Middlebrook 7H9 (BD Biosciences) supplemented with 10% BBL™ Middlebrook OADC Enrichment (BD Biosciences), 0.2% glycerol (v/v) (Sigma Aldrich), and Tween 80 (Sigma Aldrich) at 37°C, shaking. For propagation on solid agar plates, Difco™ Middlebrook 7H10 (BD Biosciences) was supplemented with 0.5% glycerol (v/v) (Sigma Aldrich). Kanamycin (20 µg/ml) was added to liquid and solid growth media where appropriate. Solid plates were incubated at 37°C for 3-5 days, unless stated otherwise.

### Mutant allele generation

C-terminal deletions were generated by PCR by deleting the region downstream of the target site. Site-directed mutations were introduced using the Q5® site-directed mutagenesis kit (New England BioLabs) as per the manufacturer’s protocol. Primers were designed using NEBaseChanger® (https://nebasechanger.neb.com/). The mutant constructs were introduced into Δ*imuA’*Δ*imuB* as per the standard electroporation protocol^15^. Kanamycin-resistant colonies were selected, and confirmed by Sanger sequencing.

### UV-induced mutagenesis and mitomycin C damage sensitivity assays

UV-induced mutagenesis assays were performed as previously described^1,2,5^ with rifampicin-resistant colonies enumerated after 5 days incubation at 37°C. For mitomycin C (MMC) damage-sensitivity assays, the cultures were grown to OD_600_∼0.4, following which a 10-fold dilution series was spotted on standard 7H10 medium and 7H10 medium supplemented with MMC. Plates were incubated for 3 days and imaged.

### Protein expression and purification

*M. smegmatis imuA’* and *imuB* were cloned into ligation independent (LIC) cloning vectors Lic1 (kanamycin resistance, N-terminal His tag) and Lic6 (streptomycin resistance, N-terminal Strep tag), respectively^16^. Co-expression of ImuB, both wild-type (His-*wt*ImuB) and mutant forms (His-ImuB_L378A, His-ImuB_V383A, and His-ImuB_L378A+V383A), with Strep-ImuA’ (and Strep-VFP-ImuA’) was performed using *E. coli* BL21(DE3) cells. Briefly, cells were grown in 2xYT medium supplemented with 1 mM magnesium sulfate, 50 μg/ml kanamycin and 50 μg/ml streptomycin. Protein expression was induced with 1 mM Isopropyl β-D-1-thiogalactopyranoside (IPTG) and cells were grown for 3 hours at 30°C. Clarified protein extracts were either loaded onto a HisTrap HP column (Cytiva Life Sciences) or directly incubated with nickel beads, and His-tagged ImuB was either with a imidazole gradient or with 500 mM imidazole, respectively. Fractions containing protein were then pooled and loaded onto StrepTrap HP columns (Cytiva Life Sciences) and eluted with 5 mM desthiobiotin. All purification steps were carried out in 50 mM Tris pH 8.5, 0.3-0.5 M NaCl, and 1 mM DTT, and samples were analysed by SDS-PAGE electrophoresis using 12% Bolt™ Bis-Tris precast gels (Invitrogen). Bands corresponding to ImuA’ in SDS-PAGE gels were quantified using the ImageLAB software from BioRad^®^.

### Size exclusion chromatography

Protein samples were injected onto a Superdex 200 PC 3.2/300 (Cytiva Life Sciences) column equilibrated in 50 mM Tris pH 8.5, 300 mM NaCl, and 1 mM DTT; thereafter, 50 μl fractions were collected.

## RESULTS

### Computational analysis of the ImuA’-ImuB interaction

Based on the experimentally determined structure of the *E. coli* RecA-ssDNA filament^17^, we first generated a homology model of the ImuA’-ImuB complex (Fig. S1). The complex included the RecA-NT homology motif (Fig. 1A) within the C-terminal region of ImuB (ImuB residues 360-387) and the ImuA’ core region (ImuA’ residues 63-194) (Fig. 1B). Using this model, as well as sequence conservation (Fig. 1A), we designed a panel of ImuB alleles containing C-terminal truncations and single amino acid substitutions that were expected to differentially affect the interaction of ImuB with ImuA’ and/or DnaE2 (Fig. 1C): (i) The first was a truncation, ImuB-ΔC380, which eliminated seven residues of the RecA-NT motif and everything downstream in the ImuB C-terminus, leaving only the α-helix (residues 360-380) predicted to be required for interaction with ImuA’; (ii) the second was also a truncation, ImuB-ΔC433, which retained the RecA-NT-like motif; (iii) the third allele mutated residues predicted to form a salt bridge including ImuA’ His99 (mutated to Asp) and ImuB Asp363 (mutated to His); and (iv) the final three alleles targeted ImuB residues – Val374, Leu378, and Val383 – for alanine substitution owing to their predicted roles in facilitating an hydrophobic interaction with ImuA’.

**Figure 1.**
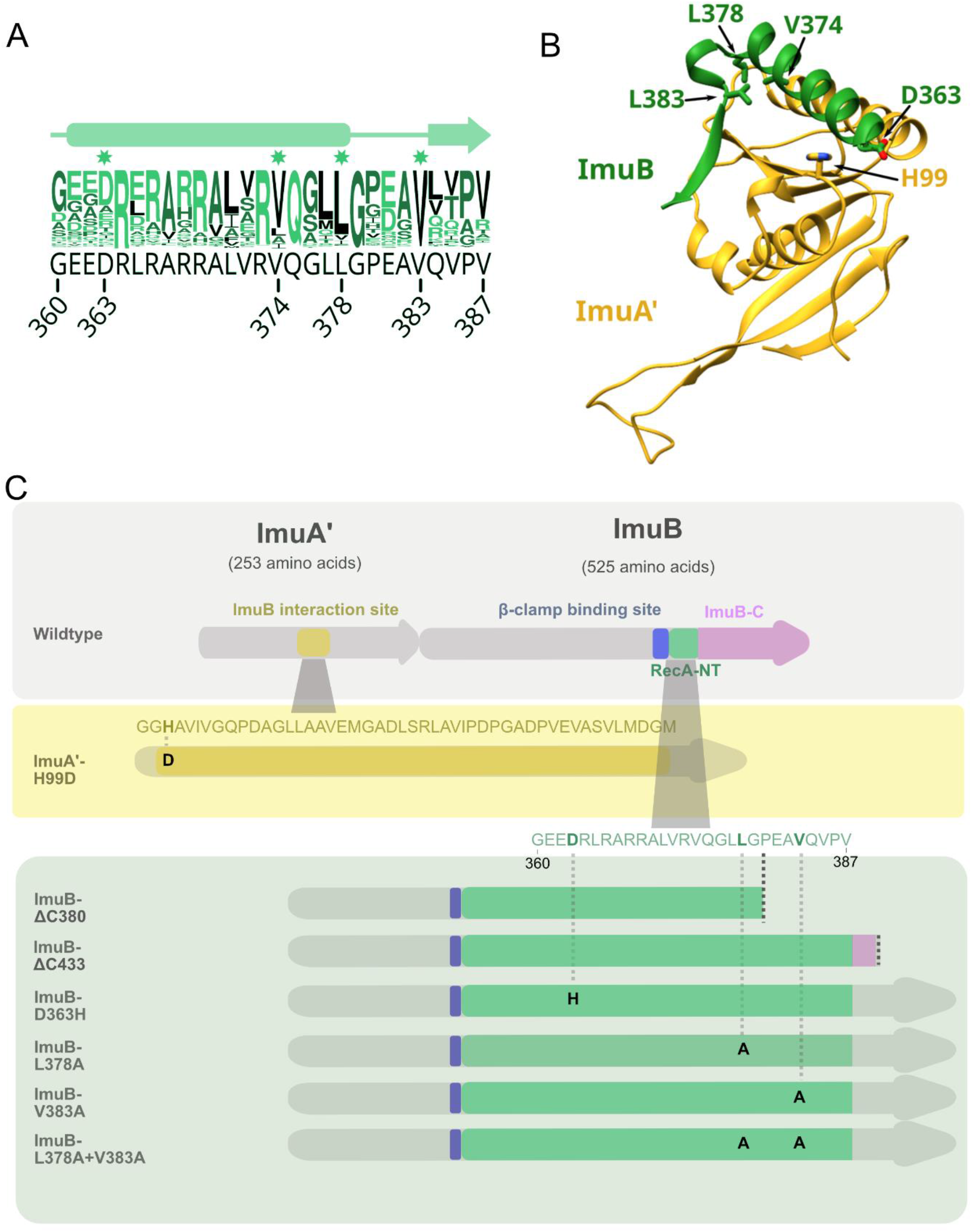
Computational model of mutasome interactions. A. Conservation logo of RecA-NT with the corresponding M. smegmatis ImuB sequence in black below. Residue positions are shown for the start and end of the motif, as well as the individual residues selected for site-directed mutagenesis. B. Interaction model between M. smegmatis ImuA’ (yellow) and ImuB (green). Displayed here are ImuA’ without the unstructured N-terminal tail (residues 1-69) in yellow and the RecA-NT-like motif of ImuB. Key residues predicted for the ImuA’-ImuB interaction are labelled and represented in sticks for both ImuA’ (H99) and ImuB (D363, V374, L378, V383). C. M. smegmatis imuA’imuB operon with predicted interaction sites. In ImuA’, the predicted ImuB interaction site stretches from G97 to M144. In ImuB, the β clamp-binding motif (Q352-W356) precedes RecA-NT (I359-V387) and is followed by ImuB-C (remaining 138 residues). The panels below indicate the site-directed mutations and ImuB C-terminal deletion mutants constructed in this study.

The *in silico* design and laboratory construction of the various mutant alleles and derivative mutants were initiated prior to the release of AlphaFold, a highly accurate protein modelling method that has transformed predictions of protein structure^11^. As soon as it became available, we used AlphaFold to generate an updated structural model of the ImuA’-ImuB complex (Fig. 2). Comparison of our original partial ImuA’-ImuB model with the AlphaFold version revealed only minor differences in the predicted interaction interface (Fig. S1); importantly, all the inferences made initially using our first homology model remained valid, allowing us to proceed with the constructs and mutants already generated. For simplicity, all analyses conducted and reported hereafter will be based on the AlphaFold model (Fig. 2).

**Figure 2.**
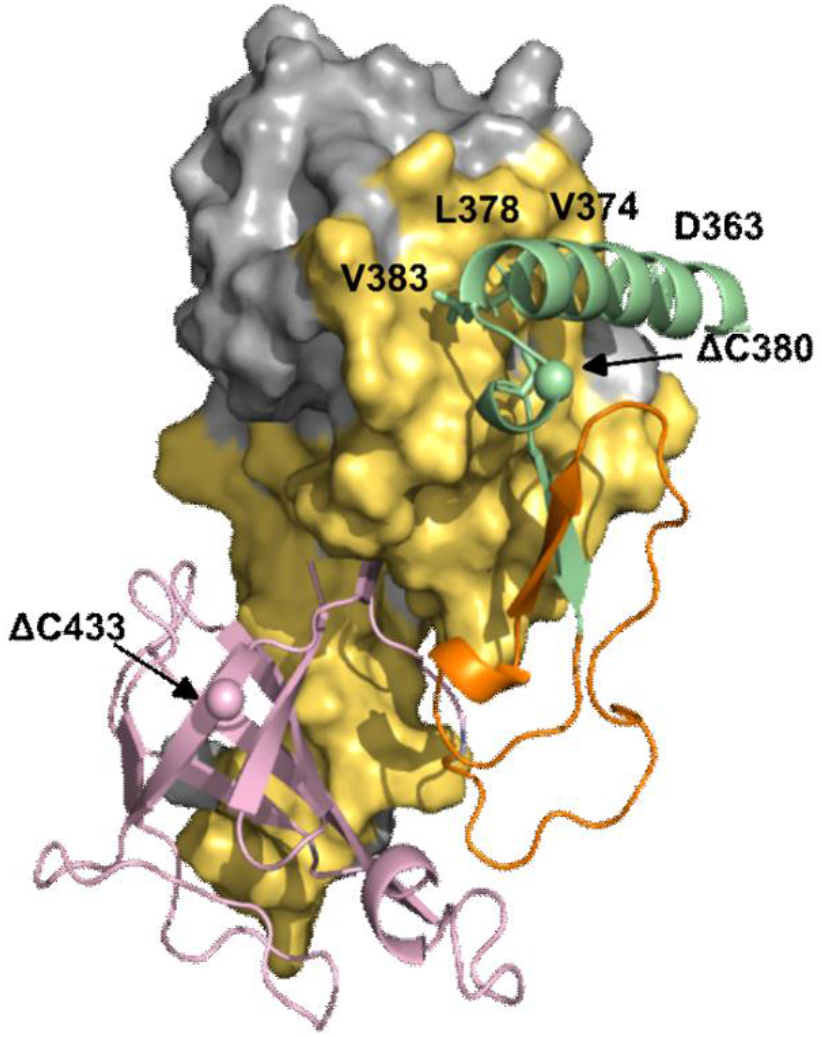
AlphaFold model of the ImuA’-ImuB complex. In yellow is the surface of ImuA’ in contact with ImuB ; green represents the RecA-NT motif (residues 360-387), pink represents the ImuB-C (residues 422-525) and orange indicates the linker between the two ImuB domains (residues 389-421). The residues where C-terminal truncations were made are highlighted with spheres.

### RecA-NT homology motif within the ImuB C-terminus is required for mutasome function

We previously reported the role of the ImuB C-terminus in interactions with DnaE1, DnaE2 and ImuA’. More specifically, we showed that the removal of the entire C-terminal region downstream of the β clamp-binding motif disrupted interactions with ImuA’ and DnaE2, abolishing mutasome function^2^. Here, we asked which region within the C-terminus of ImuB was required for mutasome function. To this end, we tested if ImuB-ΔC433, which retains RecA-NT but lacks the ImuB C-terminal region, and ImuB-ΔC380, which lacks part of the RecA-NT motif and the downstream C-terminal region, retained binding to ImuA’. The ImuB-ΔC380 deletion was incapable of complementing wildtype function in both MMC damage sensitivity (Fig. 3A) and UV mutagenesis assays (Fig 3B), suggesting disruption of mutasome assembly and/or function. To validate this inference, we shifted focus to biochemical assays utilizing purified recombinants of ImuA’ and ImuB wildtype and mutant forms. We recently showed that co-expression of ImuB and ImuA’ produces a stable complex, so that His-tagged ImuB can be used to pull down Strep-tagged ImuA’ (*via* Immobilized Metal Affinity Chromatography, IMAC, HisTrap^®^) and, reciprocally, Strep-tagged ImuA’ can pull down His-ImuB^5^ (*via* Affinity chromatography, StrepTrap^®^)^5^. To assess the contribution of ImuB-C-to the interaction with ImuA’, the two truncated versions of His-tagged ImuB – His-ImuB-ΔC380 and His-ImuB-ΔC433 – were individually co-expressed with Strep-tagged ImuA’ (Strep-ImuA’) in *E. coli* and the interaction was tested over two consecutive pull-downs (Fig. 4A, Fig. S2). Similarly to the full-length His-ImuB protein, His-ImuB-ΔC433 co-eluted with Strep-ImuA’ over the two pull-downs. However, His-ImuB-ΔC380 was unable to pull-down Strep-ImuA’ and, therefore, was absent in the subsequent pull-down using Strep-tagged ImuA’ (Fig. 4A). In combination, these results support the inferred requirement of the ImuB RecA-NT motif for functional interaction with ImuA’.

**Figure 3.**
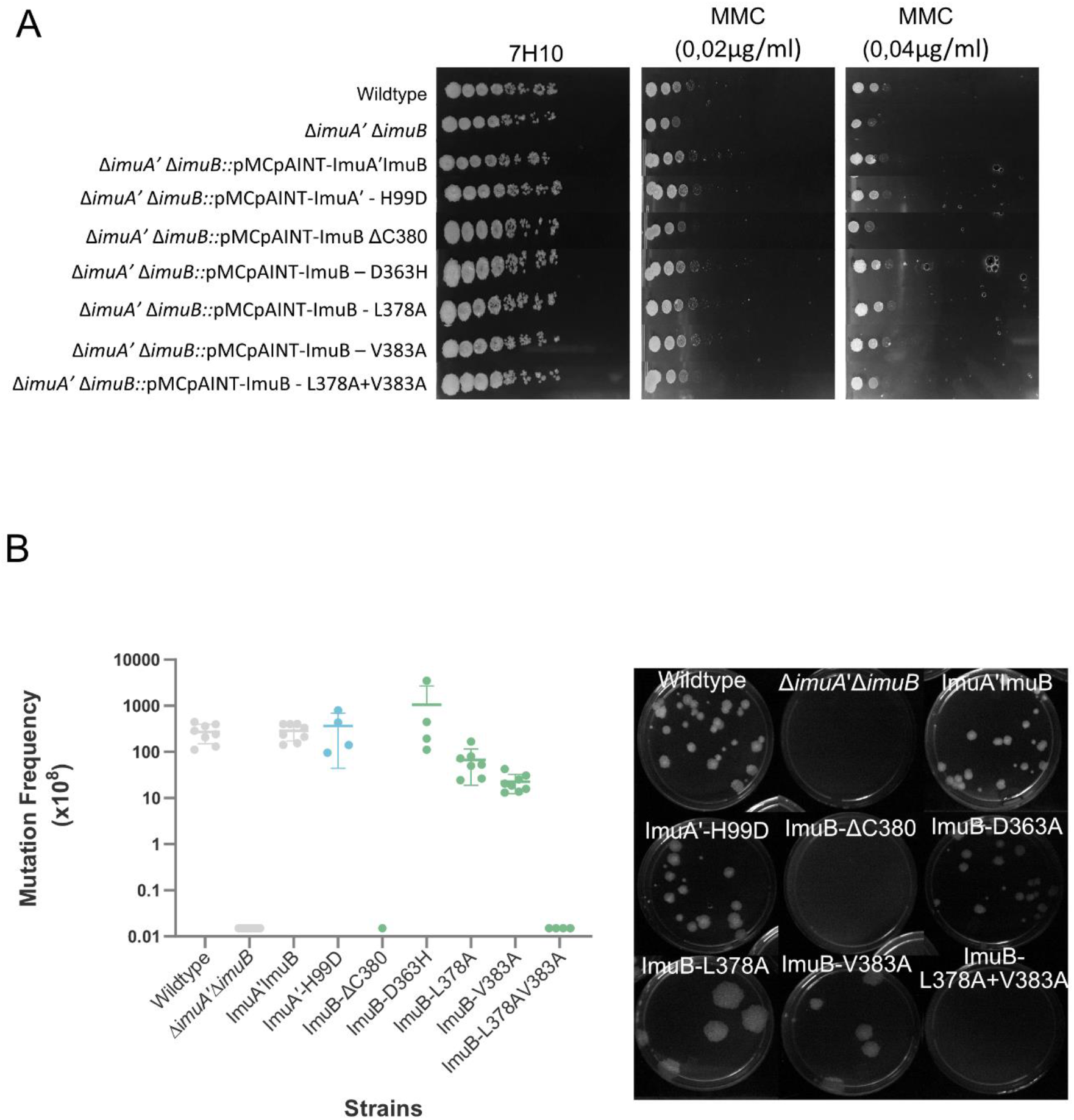
Microbiological assessment of the impact of the mutant alleles. Evaluation of functional complementation by means of A. Mitomycin C (MMC) damage sensitivity and B. UV mutagenesis assays.

**Figure 4.**
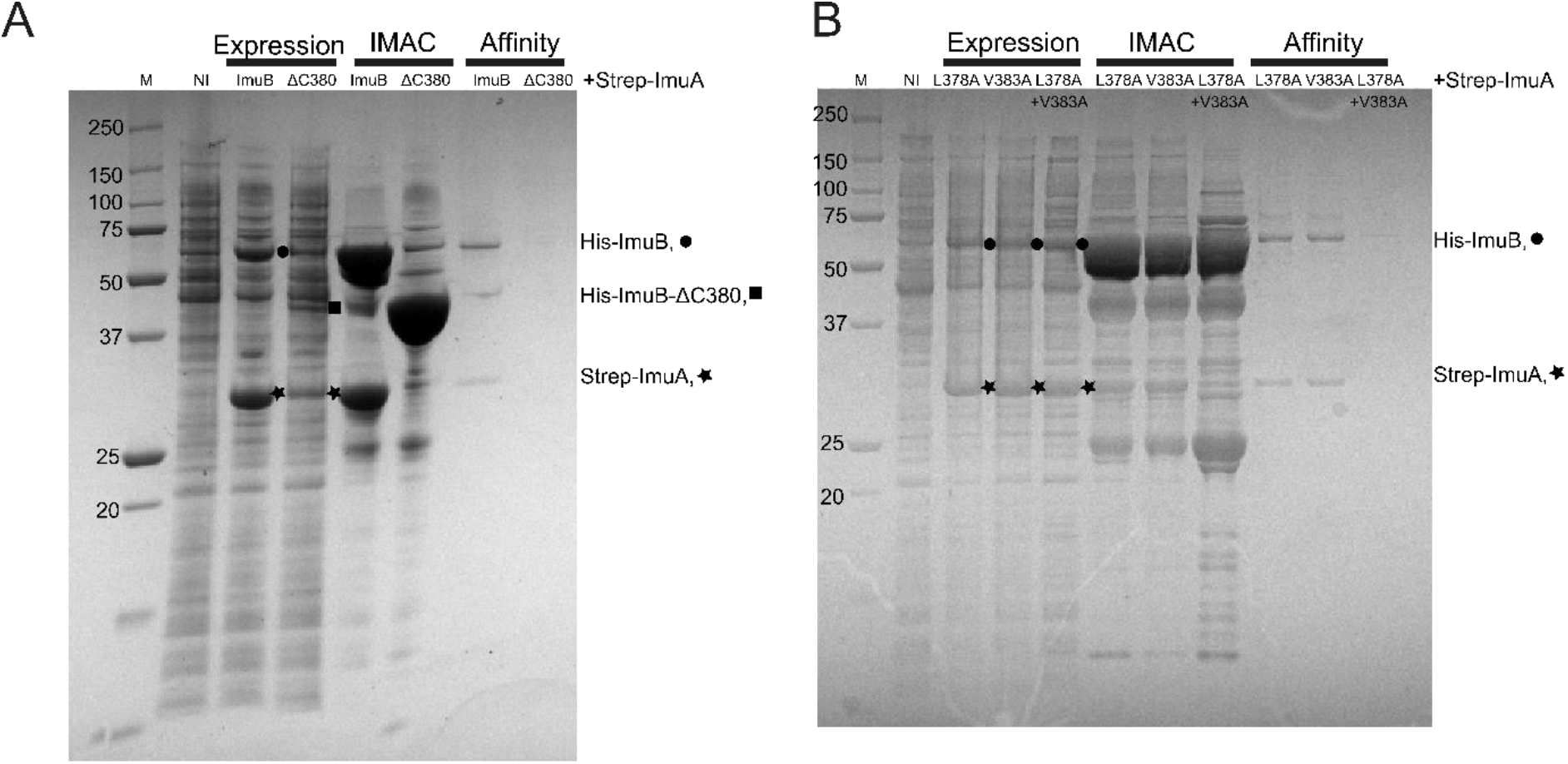
Biochemical confirmation of disruption of ImuA’-ImuB interaction in ImuBΔC380 and L378A+V383A. A. Expression samples correspond to the total expression after growth for 3h following IPTG induction. Bands corresponding to individual induced proteins are marked l (His-ImuB),□(Strep-ImuA’) and n (His-ImuBΔC380). B. SDS-PAGE analysis of the co-expression of His-ImuB mutants with Strep-ImuA’ and the two subsequent pull-downs with HisTag and StrepTag. Expression samples correspond to the total expression after growth for 3h after IPTG induction. Bands corresponding to individual induced proteins are marked l (His-ImuB mutants) and □(Strep-ImuA’).

### ImuA’-ImuB hydrophobic core interaction is required for mutasome function

To evaluate the roles of residues predicted to be at the ImuA’-ImuB interface, mutant alleles were generated and their effect on protein-protein interaction and mutasome function investigated. ImuA’ H99 and ImuB D363 – predicted to form a putative salt bridge between the two mutasome proteins – were mutated to yield ImuA’-H99D and ImuB-D363H, respectively; these mutations were expected to disrupt the electrostatic interaction in the putative salt bridge. In MMC damage sensitivity and UV mutagenesis assays, both mutant alleles restored ImuA’ and ImuB function, respectively, suggesting no significant impact on mutasome function and, by implication, ImuA’-ImuB interaction. In addition to the putative salt bridge, three ImuB amino acid residues within the RecA-NT homology motif were identified as part of the hydrophobic core of the interaction with ImuA’: V383 (start of the β-strand), L378 (end of the α-helix), and V374 (middle of the α-helix)(Fig. 1). Attempts to construct a V374A mutant failed, but V383A and L378A substitutions were generated and retained full function in the MMC damage sensitivity assays (Fig. 3A). However, these mutations appeared to cause a reproducible, albeit subtle, decrease in induced mutation frequency (Fig. 3B). Given that the effects of a single mutation might be buffered in a multi-site interaction, we decided to generate a double L378A+V383A mutant containing both amino acid substitutions. Notably, in both MMC damage sensitivity and UV mutagenesis assays, the double L378A+V383A mutant phenocopied the parental Δ*imuA*’Δ*imuB* double knockout, indicating abrogation of mutasome function.

We next assessed the impact of the L378A, V383A and L378A+V383A mutations in the ImuA’-ImuB complex formation in biochemical assays *in vitro*, adopting the approach described earlier for the C-terminal truncation of ImuB (ImuB-ΔC380). For SDS-PAGE analysis, we normalized the amount of ImuB loaded in the gel to infer the amount of ImuA’ pulled-down by each ImuB mutant. The amount of ImuA’ pulled down in the IMAC column (Histrap) by the L378A+V383A double mutant was substantially lower than the amount of ImuA’ in complex with either of the single mutants (Fig. 4B). Moreover, neither the double L378A+V383A mutant nor ImuA’ could be eluted via affinity chromatography (Affinity column) with the Strep-ImuA’ (Fig. 4B).

To improve tracking and quantification of ImuA’ during expression and purification, we used a Strep-VFP-tagged version of ImuA’ described previously^5^, allowing visualization of ImuA’ at 532 nm in the SDS-PAGE analysis. Importantly, our prior work had established that addition of the VFP fluorescence tag to the N-terminus of ImuA’ did not disrupt binding to ImuB^5^. Compared to His-ImuB, the His-ImuB-L378A+V383A protein pulled down approximately 3-fold less Strep-VFP-ImuA’, with both single mutants – His-ImuB-L378A and His-ImuB-V383A – pulling down approximately 1.5-fold less Strep-VFP-ImuA’ than His-ImuB, on average. Following size-exclusion chromatography, Strep-ImuA’ was eluted together with His-ImuB, His-ImuB-L378A, and His-ImuB-V383, confirming complex formation, albeit that the amount of ImuA’ detected for both single mutants was lower. In contrast, Strep-ImuA’ was absent in the same fraction of His-ImuB-L378A+V383A, consistent with an impaired interaction between the double ImuB L378A+V383A mutant and ImuA’. Importantly, all ImuB mutants, including the double mutant, were eluted at the same elution volume as His-ImuB after size-exclusion chromatography (Fig. S3.), indicating that the mutations introduced into ImuB did not affect protein folding or integrity. In addition, prior to all pull-down experiments, successful co-expression of Strep-ImuA’ with His-ImuB, or any of the His-ImuB mutant forms, was confirmed by the appearance of bands corresponding to the molecular weight of the target proteins after induction with IPTG (Fig. 4, Fig. S3).

## DISCUSSION

A full complement of intact, functional mutasome proteins is required for induced mutagenesis and DNA damage survival in mycobacteria^2^. Although the precise role of ImuB in mutasome function remains elusive, multiple lines of evidence support its importance in enabling key interactions with (and perhaps between) the other mutasome components, ImuA’ and DnaE2^2,5^. However, despite early evidence suggesting the importance of the ImuB C-terminus in mediating these interactions, no detail has been provided about the specific regions, motifs, or residues involved. Previous computational analyses identified two conserved regions in the C-terminus of ImuB: RecA-NT, located immediately downstream of the β clamp-binding site^7^, and ImuB-C, which is just downstream of RecA-NT^7,18^. Based on its homology to the N-terminal region of *E. coli* RecA – which is required for its oligomerisation – we hypothesized that the RecA-NT region in the C-terminus of ImuB might facilitate interaction with ImuA’ given its distant homology to RecA, its essential role in mutasome function, and the interaction between ImuA’ and ImuB inferred from Y2H studies^2^. Designing a panel of truncation and site-directed mutant ImuA’ and ImuB alleles, we have presented experimental evidence confirming the essential role of the RecA-NT motif in the ImuA’-ImuB interaction.

Based on previous observations that ImuB interacts with ImuA’ and that both ImuA’ and ImuB are required for mutasome function, we used functionality of the mutasome as a proxy for an intact ImuA’-ImuB interaction and assessed this quantitatively *in vivo* in complementation assays measuring UV-induced mutation frequencies and MMC damage sensitivities. For this purpose, we exploited a previously validated complementation system in which an integrative vector expressing the full-length *imuA’imuB* operon from its native promoter is introduced into a Δi*muA’*Δ*imuB* double deletion mutant lacking both *imuA’* and *imuB* genes^2^. Consistent with previous observations, the Δi*muA’*Δ*imuB* double knockout was hypersusceptible to MMC and showed a lower level of UV-induced mutagenesis which was restored on complementation with the wildtype *imuA’imuB* locus. In contrast, the ImuB-ΔC380 allele, which was designed to retain the key α-helix in the RecA-NT and allow at least partial interaction with ImuA’, did not restore wildtype mutasome function *in vivo* (Fig. 3). This suggests that the entire RecA-NT motif is required for a functional ImuB protein.

A limitation of these *in vivo* assays is that loss of mutasome function can’t be specifically and unequivocally ascribed to a disrupted interaction between ImuA’ and ImuB: the requirement for at least three proteins to ensure full mutasome functionality means there are potentially multiple ways in which the mutasome can fail. Therefore, to confirm the inferred disruption of the ImuA’-ImuB complex, a biochemical approach was taken. We recently reported successful expression of ImuA’ and ImuB proteins in complex^5^, noting that, in the absence of ImuB, ImuA’ appears to be mostly insoluble. We took advantage of this observation by predicting that mutants affecting the interaction between the two proteins would be expected to yield lower amounts of soluble ImuA’ rescued by co-expression with ImuB. Consistent with this interpretation, His-ImuBΔC433, in which the full RecA-NT is preserved, retained the interaction with ImuA’ (Fig. S2). In contrast, Strep-ImuA’ couldn’t be co-eluted with His-ImuBΔC380, suggesting loss of the interaction between ImuA’ and ImuB despite the retention of the key RecA-NT α-helix in the mutant ImuB allele. This observation implied that the RecA-NT α-helix alone was not enough to maintain a stable ImuA’-ImuB complex.

Disruption of the predicted salt bridge between ImuA’-H99D and ImuB-D363A did not impact the functioning of the complex *in vivo*. This suggests that either corresponding side chains are oriented away from each other in the complex or that energy contribution by this interaction is relatively small. Additionally, individual mutations in key hydrophobic residues predicted to be involved in the ImuA’-ImuB interaction (ImuB-L378A and ImuB-V383A) resulted in a very small but reproducible drop in the induced mutagenesis frequency (Fig. 3B) despite having no visible effect on MMC damage sensitivity (Fig. 3A). However, an ImuB-L378A+V383A double mutant showed complete loss of function in both assays (Fig. 3). These observations were recapitulated in biochemical assays in which His-ImuB-L378A and His-ImuB-V383A recovered slightly lower levels of Strep-ImuA’ than His-ImuB, while His-ImuB-L378A+V383A scarcely recovered any Strep-ImuA’ (Fig. 4B). In combination, these results suggest that altering the residues individually is not enough to disrupt the interaction but, together, these residues are vital in maintaining a stable ImuA’-ImuB interaction. We also confirmed that substitution of the residues did not alter the protein elution volume by size-exclusion chromatography and therefore did not have a significant impact in the folding nor in the integrity of ImuB (Fig S2).

The RecA-NT motif was identified based on homology to the region in *E. coli* RecA responsible for oligomerisation by binding an upstream RecA molecule on the RecA filament^7^. The results presented here reveal that the RecA-NT motif also facilitates binding with ImuA’, a distant homolog of RecA. It is tempting to suggest analogy with the interaction between the RecA-NT motif in the multicomplex *E. coli* multi-protein complex mutasome (UmuD’_2_C-RecA-ATP), which is absent in mycobacteria. Whether or not ATP is needed for the regulation of ImuA’B-DnaE2 activity is still not clear.

A predicted structure of the ImuA’B complex was generated using AlphaFold (Fig. 2). From this model, the ImuA’-ImuB interaction interface appears more extensive than the RecA-RecA interface (Fig. 2, Table S1). The C-terminal region of ImuB beyond RecA-NT (ImuB-C) is also predicted to be involved in binding ImuA’, contributing similar surface area to the interaction as that of RecA-NT (Table S1). Nonetheless, the ImuB RecA-NT-like motif appears to be the key element for binding ImuA’, because disrupting the hydrophobic residues in RecA-NT almost completely abolished ImuA’-ImuB interaction (Fig. 4B). In further support of this conclusion, an ImuB-ΔC433 truncation, which preserves RecA-NT but eliminates ImuB-C, retained ImuA’ binding ability in biochemical assays (Fig. S2). The potential importance of the mycobacterial mutasome – minimally comprising ImuA’, ImuB, and DnaE2 – in *M. tuberculosis* evolution and acquisition of drug resistance has been recognized previously ^1,2,19,20^. We are optimistic the results presented will allow for a better understanding of the assembly and operation of this mutagenic machinery, and will inform ongoing efforts towards developing novel “anti-evolution” drugs^21–24^ for *M. tuberculosis* and other organisms encoding homologous systems.

## SUPPLEMENTARY DATA

Supplementary Material is available online.

## ACKNOWLEDGEMENT

The authors gratefully acknowledge the facility and expertise of the Protein Facility at the Cell and Chemical Biology Department (Leiden University Medical Center, Leiden, The Netherlands) and are particularly grateful to Pedro J. B. Pereira (Instituto de Biologia Molecular e Celular, Porto, Portugal) for insightful comments on the manuscript.

