## Supplementary Material for "RecA-NT homology motif in ImuB is essential for mycobacterial ImuA’-ImuB protein interaction and mutasome function"

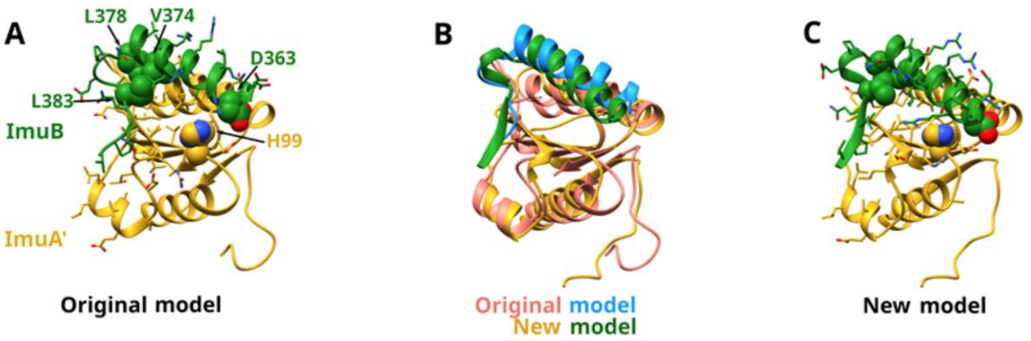


***Fig. S1*. Comparison of original Modeller model A.** *to new AlphaFold model* ***C.*** *with* ***B.*** *showing both models superimposed (orginal model – salmon/blue, new model - gold/green). Only structurally corresponding parts to the original model are shown for the new model: 63-178 of ImuA' and 360-387 of ImuB. The original model includes 67-194 of ImuA', but the C-terminal beta-strand is modeled incorrectly according to the AlphaFold prediction.*

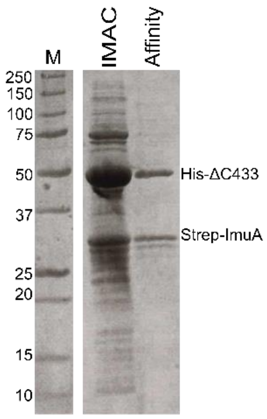


***Fig. S2 - His-ImuBΔC433, in which the full RecA-NT is preserved, forms a stable complex with ImuA'.*** SDS-PAGE analysis of the purification of His-ImuBΔ433 with Strep-ImuA' after two sequential pull-downs with HisTag and StrepTag, respectively.

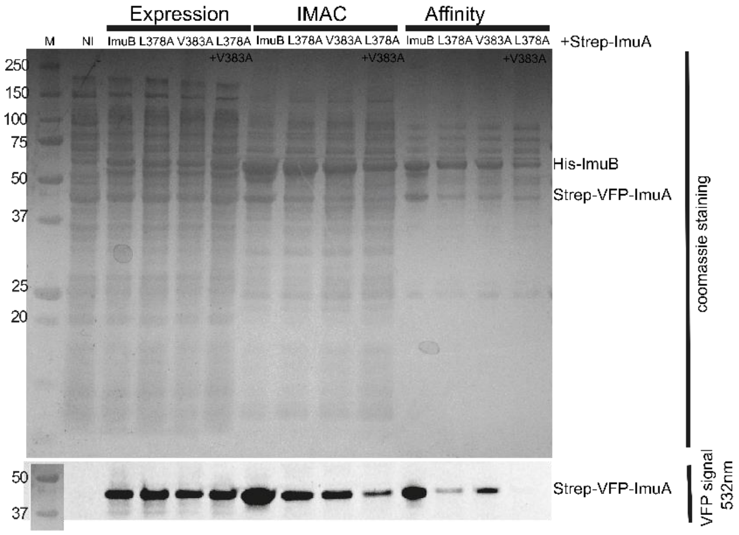


***Fig. S3. Biochemical confirmation of disruption of ImuA'-ImuB interaction in L378A+V383A (double mutant, DM) using a fluorescent tagged version of ImuA´(Strep-VFP-ImuA´).*** *SDS-PAGE analysis of the co-expression of His-ImuB mutants with Strep-ImuA' and the two subsequent pull-downs, followed by size-exclusion chromatography. Expression samples correspond to the total expression after growth for 3h following induction with IPTG. Above is the SDS-PAGE gel stained with Coomassie and bellow is the same gel imaged at 532nm for detection of the VFP fluorescent tag in ImuA'.*

***Table S1 Interface surface areas between ImuB constructs and full length ImuA'****. Calculated with VoroContacts from the ImuA'-ImuB AlphaFold model.*

| **ImuB construct** | **Interface area with ImuA', Å$^2$** |
|---|---|
| ImuB, full | 1804 |
| ImuB, 360-388 (RecA-NT motif) | 694 |
| ImuB, 422-525 (ImuB-C) | 960 |
| ImuB, 1-359 (C-terminal region removed) | 0 |
